# Fast-SG: An alignment-free algorithm for hybrid assembly

**DOI:** 10.1101/209122

**Authors:** Alex Di Genova, Gonzalo A. Ruz, Marie-France Sagot, Alejandro Maass

## Abstract

Long read sequencing technologies are the ultimate solution for genome repeats, allowing near reference level reconstructions of large genomes. However, long read *de novo* assembly pipelines are computationally intense and require a considerable amount of coverage, thereby hindering their broad application to the assembly of large genomes. Alternatively, hybrid assembly methods which combine short and long read sequencing technologies can reduce the time and cost required to produce *de novo* assemblies of large genomes. In this paper, we propose a new method, called FAST-SG, which uses a new ultra-fast alignment-free algorithm specifically designed for constructing a scaffolding graph using light-weight data structures. FAST-SG can construct the graph from either short or long reads. This allows the reuse of efficient algorithms designed for short read data and permits the definition of novel modular hybrid assembly pipelines. Using comprehensive standard datasets and benchmarks, we show how FAST-SG outperforms the state-of-the-art short read aligners when building the scaffolding graph, and can be used to extract linking information from either raw or error-corrected long reads. We also show how a hybrid assembly approach using FAST-SG with shallow long read coverage (5X) and moderate computational resources can produce long-range and accurate reconstructions of the genomes of *Arabidopsis thaliana* (Ler-0) and human (NA12878).

## Introduction

The major challenge of whole genome *de novo* assembly is to solve repeats (1, 2). These correspond to nearly identical genomic sequences that occur at multiple locations in a genome. To address this challenge, two major types of approaches have been proposed, one using paired short reads (3) and the other long reads (4).

In the second case, the aim is to hopefully entirely capture the repeats within the long reads. The non repeated suffix and prefix sequences of such long reads are used to compute unique overlaps, which then allow to unambiguously expand the original reads into larger ones, called contigs, in a process that may sometimes (but not always) directly lead to inferring the entire genomic sequence.

The first type of approach instead needs to be associated to an operation called genome scaffolding. The short reads are still first assembled into contigs as above, either by also computing overlaps (5) or by using de Bruijn graphs (6). The contigs obtained in this case will however not span the whole genome. Indeed, most often they will be much shorter. They then need to be joined (i.e. linked together) in a second step. The linking information is in general provided by paired-end or mate-pair sequencing. Commonly, genomic fragments larger than 1kb from which both ends are sequenced are denoted as mate-pair libraries, otherwise they are referred to in the literature as pair-end libraries. Genome scaffolding that uses paired short reads introduce gaps (i.e. unknown sequences) between the contigs, thereby once again not leading to the entire genomic sequence but to a set of so-called scaffold sequences, or scaffolds for short. A scaffold thus represents a set of ordered and oriented contigs.

The genome scaffolding problem was first formulated by Huson *et al.*(7). The method proposed by the authors started by building what is called a scaffolding graph where the nodes represent the contigs and the edges encode the number of mate-pairs (weight), the orientation and the distance between two different contigs. A greedy algorithm is then used to heuristically obtain optimal paths that will correspond to the scaffold sequences.

Most of the scaffolding methods that have been developed since use the same type of graph, built with ultra-fast short-read aligners (8, 9, 10) as a foundation for the scaffolding (3). Algorithmic innovations in the area are mainly focused on how to select optimal paths (usually those of maximal weight) and thus obtain large and accurate scaffolds. Various approaches have been proposed, based on dynamic programing (11), breadth-first search (12), maximum weight matching (13), or branch and bound (14), among others.

The new long read sequencing technologies (Pacific Biosciences, Oxford Nanopore) suddenly changed the genome assembly scene by producing very long (*>*10kb) reads that however contain a high level of errors (on average 15% at the current time). These new technologies nevertheless extended the landscape of solvable repeat sequences (15). Currently, *de novo* assemblers that use such long reads (4, 16) are thus able to finish bacterial genomes and to produce highly continuous reconstructions of human genomes (4). However, *de novo* assemblies of large genomes based on computing overlaps (5) are computationally intense (4) and require a considerable amount of coverage (*>*30X) to error-correct the inaccurate long read sequences by self-correction methods, thereby hindering a broad application of these methods to the *de novo* assembly of large genomes (BioRxiv: https://doi.org/10.1101/128835).

*de novo* assemblies using long reads have nevertheless proven to be scalable to chromosomes (17, 18) when associated with complementary long range information from novel library preparation techniques (19, 20). Such new experimental libraries are sequenced on Illumina machines leading to conventional paired-end reads. DOVETAIL genomics (19) thus produces useful linking information in the range of 1-200kb, while 10X genomics (21) produces, by using barcodes in a clever manner, linked-reads in the range of up to 100kb. Both technologies then use such long-range information within their assembly pipelines (19, 21) to build a scaffolding graph to which they apply their own algorithmic solutions to obtain the scaffold sequences. Both technologies were conceived with the aim of replacing the expensive and time consuming experimental protocols required to produce long-range mate-pair libraries (22, 23) with short-read sequencing.

In principle, long-range information can be extracted directly from long reads in ranges restricted to the latter’s actual sizes. Such information can then be used to devise a hybrid assembly method, where high quality contigs from short read assemblies are used as nodes of the scaffolding graph, edges are created using linking information from the long reads, and the scaffolds are generated by a short read scaffolder. However, there is currently a lack of algorithms for building a scaffolding graph from the long reads. Such algorithm would allow the reuse of efficient existing short-read algorithms to compose novel hybrid assembly pipelines.

Being able to (i) build such a graph from either short or long reads in an ultra-fast way, with moderate computational resources, while (ii) keeping the structure standard enough to be compatible with the existing efficient short read scaffolders are the main challenges that we address in this paper. The method that we propose, called FAST-SG, uses an alignment-free algorithm (24) strategy as well as information from varied sequence sources (Illumina, Pacific Biosciences and Oxford Nanopore), and was conceived to maximize scalability, speed, and modularity. The latter characteristic in particular allows to define novel hybrid assembly pipelines, which permits the efficient assembly of large genomes.

FAST-SG was extensively tested using a comprehensive set of standard datasets (3, 25). Four benchmarks were performed. The first had for objective comparing FAST-SG to the state-of-the-art short read aligners for building the scaffolding graph.

In the second benchmark, the capacity of FAST-SG to extract linking information from long read technologies is shown, and we then tested how short read scaffolders fed with FAST-SG compare against a dedicated long read scaffolder. The third and fourth benchmarks explore what is the major result of this paper, namely the full hybrid genome assembly solutions that can be obtained using FAST-SG as a component. We start by analyzing the amount of long read coverage required for such hybrid solutions and compare the performance obtained with our approach against a state-of-the-art long read assembly pipeline (4) when building the genome of *Arabidopsis thaliana* (Ler-0). In the final benchmark, we then perform the hybrid assembly of a complete human genome (NA12878). We first show that FAST-SG can extract even Bacterial Artificial Chromosome (BAC) clone size (180kb) linking information from ultra-long Nanopore reads (BioRxiv: https://doi.org/10.1101/128835) at low coverage (5X), without error correction and using moderate computational resources. We then compare the results we obtain with those produced by *de novo* long-read assembly pipelines and third generation mapping technologies. We conclude by discussing how the modular hybrid assembly approach with FAST-SG as a main component that we propose in this paper can be extended to use long reads to fill the gaps and error-correct the scaffold sequences.

## Materials and Methods

### FAST-SG index

The FAST-SG index consists of all the unique *k*-mers present in the set of target contigs at a given *k*-mer length. For each of them, we store the position, the strand and the contig of origin, using lightweight data structures such as Minimal Perfect Hashing (ArXiv: https://arxiv.org/abs/1702.03154) and Probabilistic Dictionary (ArXiv: https://arxiv.org/abs/1703.00667). In a first step, we define the unique *k*-mers as being those having a frequency equal to 1 from the total set of distinct *k*-mers present in the target contig/genome sequences. To identify unique *k*-mers, we use KMC3 (26), an ultrafast, parallel and memory frugal *k*-mer counter. In a second step, each unique *k*-mer is hashed to the space of [2^0^,2^64^] using a rolling hash function (27) and with hash values written on the fly to a binary file. Rolling hashing has the nice property of computing hash values for consecutive *k*-mers in a sequence in 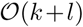 time, where *k* is the *k*-mer length; *l* is the sequence length and *k < l*. We use an efficient library implementation of rolling hash algorithms called NTHASH (28), which implements a barrel shift function and a seed table of integers to compute hash values in both DNA strands faster.

In a third step, the static hash values stored in the binary file are used as input to create a Minimal Perfect Hash Function (henceforth denoted by MPHF). MPHF provides a collision-free and minimal space way to store and look-up hash values in constant worst-case access time for static sets. We use the library implementation provided by Limasset *et al.* (ArXiv: https://arxiv.org/abs/1702.03154), called BBHASH, which is simple, parallel, fast and memory frugal. Moreover, it can store 10^10^ hash values using moderate computational resources (5Gb). The major feature of MPHF is its ability to map each key of *S* (in our case, the unique *k*-mer hashed values) to an integer in the interval [1*,N*] (injective function), with *N* = *|S|*, while avoiding the implicit storage of hash values by using cascade hash functions in conjunction with bit vectors. A significant parameter of BBHASH is the *γ* (gamma) factor. We use a *γ* factor equal to 4, which is an optimal value for fast query time, fast construction and low memory usage. When performing a query in the MPHF structure, it returns an index in the interval of [1*,N*] which has the same size of the static set *S*, allowing to store related data for each *s ∈ S* using simple arrays. If we query a key not present in the initial static set *S*, MPHF could return a value in the interval [1*,N*] that is a false positive (ArXiv: https://arxiv.org/abs/1703.00667).

In a fourth step, to control the false positive rate (*p*) of MPHF, we use a probabilistic set (ArXiv: https://arxiv.org/abs/1703.00667). For each indexed element *s ∈ S* (unique *k*-mers), we store a fingerprint value using 16 bits in an array of size *N* = *|S|* at the corresponding MPHF index of *s*. The fingerprint is built by re-hashing the hash value of *s* using the xor-shift hash function in the range [2^0^,2^16^] and storing it in a bit-set array structure. We selected a fingerprint of size 16 bits, because it has a low false positive rate *p* = 1/2^16^ = 0.0000152.

Finally, we added the associated contig id, strand and coordinate values of each unique *k*-mer stored in the MPHF and the probabilistic dictionary (MPHF-PD), by performing a single pass through the set of contigs/genome sequences, using the same *k*-mer size. For each *k*-mer hit, we store the values (contig id, coordinate and strand) in the index returned by the MPHF-PD structure using three vectors having the same size as the set *S*. After storing all the associated values, we end our index construction and return a reference to the new object. This object is the FAST-SG index. The memory required per *k*-mer is composed of 6 bits for the MPHF, 16 bits for the probabilistic dictionary, 32 bits for the contig id, 32 bits for the position, and 1 bit for the strand, adding to a total memory of 87 bits.

### FAST-SG alignment-free method

The core of FAST-SG is an alignment-free algorithm specifically designed to construct the scaffolding graph from either short or long reads using light-weight data structures. Such graphs are built using as information the read pairs that map uniquely to different contigs. If the mappings are within an expected distance from one another given the respective orientation of the reads, an edge is added to the graph between the contigs (3). The uniqueness property of the mapping is ensured by its high quality score which represents the confidence that the read indeed belongs to the reported genomic location (9, 10). When a read belongs to two possible genomic locations, a score of 0 is commonly assigned.

Current short read aligners identify the high quality score mappings by indexing all the *k*-mers present in the set of contigs and using a seed-and-extend (9, 10) alignment approach. Instead, in FAST-SG, only the *k*-mers having a frequency equal to 1 are considered and no alignment is performed. After building the FAST-SG index, the contig location for a pair of reads is determined following a number of steps as illustrated in Figure 1a.

**Figure 1.**
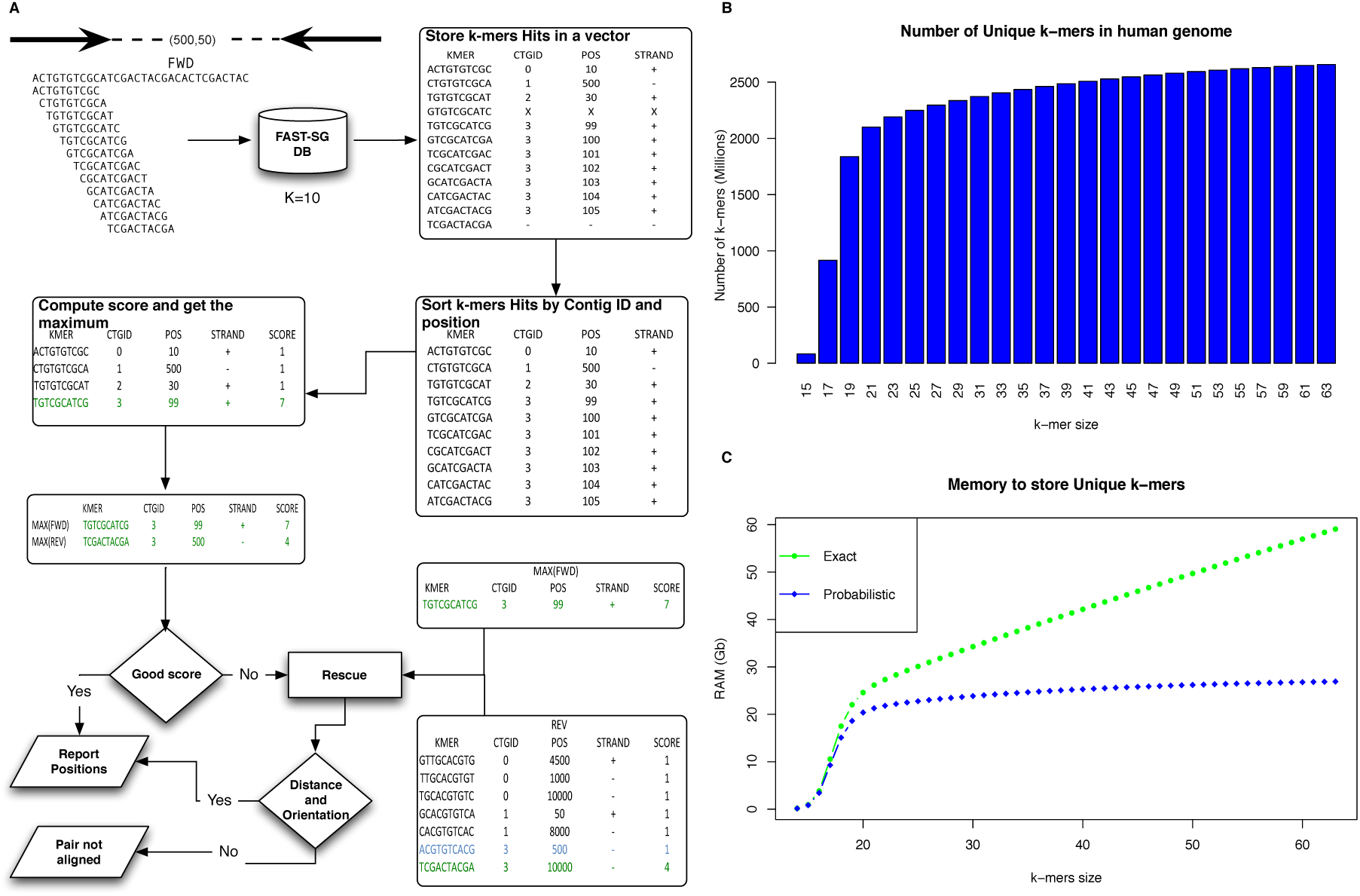
A) Overview of the F_AST_-SG algorithm. B) Number of unique *k*-mers (y-axis) in the human genome GRCh38.p10 as a function of the *k*-mer size (x-axis). C) Memory required for indexing the unique *k*-mers of the human genome by F_AST_-SG and using an exact implementation. The blue dotted-line shows the memory required by F_AST_-SG as a function of the *k*-mer size. In green is shown the memory required by an exact implementation which uses two bits per base. The amount of memory used by such implementation increases as a function of the *k*-mer size (x-axis). The memory of the index used in F_AST_-SG only increases with the number of *k*-mers to store.

The first step performs look-ups of the *k*-mers of the forward (resp. reverse) read sequence (on both strands by using a rolling hash function) in the FAST-SG index, and fills a vector of hits of a predefined size. The size of the vector depends on the error rate of the sequencing technology. The default chosen in FAST-SG is of 10 for Illumina and 20 for the long-read technologies. In a second step, the forward (resp. reverse) vector of the *k*-mer hits is sorted by contig and, inside each contig, by coordinate. In the third step, a score is computed for the forward (resp. reverse) read that corresponds to the maximum number of hits falling inside a window of size equal to the length of the read. If the score of both reads in a pair reaches a predefined minimum, in a fourth step the genomic location of the pair is reported. Otherwise, a pair rescue is attempted (fifth step) by fixing the location of the best scored read and looking for a *k*-mer hit in the mate pair that satisfies the expected distance and orientation (Figure 1a).

A major parameter of the algorithm is the *k*-mer size as this governs the number of unique *k*-mers to be indexed in a given genome, or in our case, a set of contigs. In Figure 1b, we show how the number of unique *k*-mers increases as a function of the *k*-mer size in the human genome (GRCh38.p10). However, large *k*-mers need reads with low error rates for a successful match. To define an appropriate *k*-mer size, it is necessary to take into account both the error rate and the length of the query sequence. Almost all short read aligners use as seeds short *k*-mers (15-32 base pairs) because they have a low probability of containing errors and provide enough specificity (9, 10, 29). Additionally, the available long read algorithms such as CANU (4), LORDEC (30), and MASURCA (31) among others, employ short *k*-mers (15-19 base pairs) at some stages to deal with the large error rates (15%) present in the current long read technologies. In practice, FAST-SG supports a *k*-mer size of up to 256 base pairs, but for the Illumina reads, values of k between 15 and 80 were tested while for long reads, these ranged from 15 to 22 base pairs (due to the high base-error rate present in the current long read technologies), which according to our benchmarks provide enough specificity, even for large genomes (Figure 1b). There are for instance 1.83 billion unique 19-mers (Figure 1b) in the human genome, which is a good approximation of the non-repetitive regions for this genome (2).

Another issue of working with *k*-mers is the memory required for storing them for fast look-ups. This was addressed by implementing a novel probabilistic data structure (FAST-SG index) which only requires 87 bits per *k*-mer, while memory increases as a function of the number of unique *k*-mers to store (Figure 1c). To index in memory all the unique *k*-mers of the human genome at a given *k*-mer size (<256bp) therefore requires less than 30Gb of memory (Figure 1c).

Finally, the genomic location of the read pairs is reported using a single representative unique *k*-mer for each read in SAM format (32), thus allowing for an easy integration with scaffolders that support this standard format. The steps of scoring and pair rescuing follow some of the ideas used in the SSAHA (29) and BWA-MEM (ArXiv: https://arxiv.org/abs/1303.3997) aligners.

### Illumina mate-pair reads alignment

Illumina mate-pair reads are aligned using the algorithm described previously (FAST-SG alignment-free strategy). The forward read (QF) is iterated *k*-mer by *k*-mer where for each *k*-mer, we ask if it is present in the FAST-SG index until 10 hits are stored in the vector *vectorFUH*. If the score of QF is larger than 3, we attempt to fill the vector *vectorRUH* (QR) of the reverse read. Then, if the score of each read is larger than 5, the positions are reported. Otherwise, we attempt pair rescue by fixing the position of the best-scored read and requiring a minimum score of 4 for the rescued read. These parameters of minimum and pair-rescue scores were set from empirically derived defaults (Illumina alignment benchmark). Such default short read parameters can be modified by the user.

### Extraction of synthetic pairs from long reads

Synthetic pairs of reads (QF and QR) are extracted from the long-read sequences having a default read length of 200 base pairs in forward-reverse orientation and separated by a distance D (insert size). Multiples values of D can be specified to comprehensively extract linking information from the long reads. After extracting a synthetic pair, each query sequence (QF and QR) is aligned using the algorithm described previously (FAST-SG alignment-free strategy). A minimum score of 15 and a minimum rescue score of 4 are used as default parameters. Then, as default, a moving window of 100bp is adopted to extract another pair, until the complete long read sequence is scanned. The default long read parameters can be modified by the user.

### Estimation of the genomic library parameters

The genomic library parameters for insert size, standard deviation and orientation are estimated using a subset of the mate-pair sequences in order to use them in the rescue step of FAST-SG. These subsets of mate-pair reads are aligned to the target contigs/genomes, and the read pairs located within contigs are used to estimate the library parameters. For Illumina, we use a total of 100,000 pairs which are aligned to the target sequences using a minimum score of 8 and without pair rescue. Then, for each aligned pair within contigs, we save the pair orientation and distance. To infer the average insert size and standard deviation, we remove 10 percent outliers from both tails of the values stored by sorting the observed insert sizes by increasing order. The orientation is computed using a majority rule on the four possible orientations for a pair of reads (FR, RR, FF, RF). For long reads, we use a total of 1,000 long read sequences and we extract the specified insert sizes to infer the average insert size and standard deviation as for the Illumina libraries. The orientation for the synthetic libraries is not estimated because all pairs are created in forward-reverse orientation.

### Concurrent steps of FAST-SG

The index construction and alignment steps in FAST-SG are concurrent. The FAST-SG index can use multiple threads to construct the MPHF (ArXiv: https://arxiv.org/abs/1702.03154) and store the associated *k*-mer information (contig id, coordinate, strand). Chunks of 5Mb of contig sequences are used to populate in parallel the FAST-SG index. The FAST-SG alignment step is concurrent by taking chunks of 500,000 and 1,000 for the short and long reads respectively. The concurrent steps are implemented using the PTHREAD library. The user specifies the number of CPUs to be used.

### Datasets and software

We collected a comprehensive collection of standard datasets (Table 1) which are frequently used to benchmark the new sequencing technologies, scaffolding tools or genome assembly pipelines.

**Table 1.** Sequencing datasets and Illumina assemblies used to evaluate the performance of F_AST_-SG

| Short read datasets |  |  |  |  |  |  |
| --- | --- | --- | --- | --- | --- | --- |
|  | #Reads | Read length | Insert size | SRA/ENA | Illumina assemblies |  |
|  |  |  |  |  | #Contigs | N50 |
| *S.aureus* | 3,494,070 | 37 | 3,500 | SRR022865 | 170 | 47,016 |
| *R.sphaeroides* | 2,050,868 | 101 | 3,500 | SRR034528 | 577 | 15,351 |
| *P.falciparum* (short) | 52,542,302 | 76 | 550 | ERR034295 | 9,318 | 2,995 |
| *P.falciparum* (long) | 1,562,080 | 75 | 3,000 | ERR163027 |  |  |
| *H.sapiens* chr14 | 22,669,408 | 101 | 2,600 | SRR067771 | 19,936 | 12,963 |
| Long read datasets |  |  |  |  |  |  |
|  | #Reads | Average read length (bp) | Technology | Machine | Illumina assemblies |  |
|  |  |  |  |  | #Contigs | N50 |
| *E.coli* K12 | 164,472 | 9,009 | ONT | R9.2 | 140 | 106,241 |
|  | 1,192,955 | 4,412 | PacBio | Sequel System |  |  |
|  | 22,391,084 | 298 | Illumina | Myseq |  |  |
| *S.cerevisiae* W303 | 594,243 | 4,795 | PacBio | PacBio | 890 | 52,324 |
| *A. thaliana* (Ler-0) | 561,176 | 9,633 | PacBio | Sequel System | 2,384 | 320,571 |
|  | 46,129,480 | 300 | Illumina | Myseq |  |  |
| Human (NA12878) | 1,415,868 | 16,324 | ONT | R9.4 | 37,393 | 202,174 |

Further details are provided in the Methods Section and in the Supplementary Material 1.

In particular, to assess the performance of FAST-SG for constructing the scaffolding graph from short reads, we employed all the short read datasets and Illumina assemblies defined in Hunt *et al.* (3). These short read datasets include the genomes of *Staphylococcus aureus*, *Rhodobacter sphaeroides*, *Plasmodium falciparum* and the human chromosome 14 (Table 1), and are commonly used as the gold standard for validation of the scaffolding tools (11, 12, 13, 14).

Long read datasets were then used to investigate the capacity of FAST-SG to extract linking information from long reads, and then the performance of short read scaffolders fed with FAST-SG when compared to a dedicated long read scaffolder. In the first case, the genome of *Escherichia coli K.12* was adopted as it has been sequenced by multiple long read technologies and is commonly used to validate the long read algorithms (4). In the second case, both *E. coli K12* and *Saccharomices cerevisiae* W303 (Table 1) were employed to prove that short read scaffolders can be transformed into long read ones.

To explore the amount of long read coverage required by the hybrid solutions we propose and compare the performance of the latter to the results obtained by CANU (4), a state-of-the-art long read assembler, we used in a first step the genome of *Arabidopsis thaliana*, and then in a second step, a complete human genome (NA12878, Table 1). NA12878 was selected because it have been sequenced on a variety of platforms (19, 21, 33) and as well as assembled by a variety of algorithms (4, 19, 21, 31). It thus allows to compare the complete landscape of currently available long-range technologies and assembly pipelines. Finally, All software and the reference genomes used are described in the Supplementary Material 1.

### Short and long reads benchmarks

The Illumina alignment benchmark was performed as described in the Supplementary Material 2. The Illumina scaffolding benchmarks are detailed in the Supplementary Material 3. The long read scaffolding benchmarks are described in the Supplementary Material 4. The hybrid scaffolding benchmarks are described in the Supplementary Materials 4 and 5. All scaffold sequences generated from alignments produced by FAST-SG or together with the short read aligners or and by LINKS were evaluated following the standard defined by Hunt *et al.* (3). For each dataset, the true contig layout is known and the scaffold sequences were compared against it in order to determine the following scaffolding errors (represented as a bitwise flag):

0 = Correct pair of contigs.

1 = Contigs originated from same reference sequence, but their orientation in the scaffolds is incorrect. 2 = Contigs originated from different reference sequences.

4 = Contigs originated from the same reference sequence, but are the wrong distance apart.

5 = 4+1, Contigs originated from same reference sequence, but their orientation and distance in the scaffold is incorrect. 8 = Contigs originated from the same reference sequence, but are not in the correct order.

12 = 8+4, Contigs originated from the same reference sequence, but are not in the correct order and distance.

From the previous values, we computed the F-Score metric, which was first introduced by Mandric and Zelikovsky (13), and adopted in Luo *et al.* (14) also with the purpose of improving and summarising in a single metric the performance of a scaffolding tool. In brief, if we denote by P the number of potential joins that can be made, TP the number of correct joins performed by a scaffolder (true positives), and FP the number of wrong joins (false positives), we can calculate the following quality metrics:

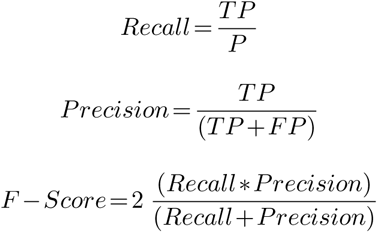

The structural quality of the hybrid and *de novo* assemblies was determined via direct comparison against the nearest reference genomes available using NUCMER (34) and reported using the GAGE statistics (25) which from 1-to-1 alignments evaluates both the identity and the structural breakpoints (inversions, relocations and translocations). All commands executed in each benchmark are specified in the Supplementary Materials 2 to 4.

## Results

### Comparison of FAST-SG with the state-of-the-art short read aligners

We assessed the performance of FAST-SG for aligning short reads on simulated Illumina data from the complete human reference genome (GRCh38.p10, see Table 2 and Supplementary Material 2) together with BOWTIE (8), BOWTIE2 (10), BWA-MEM (ArXiv: https://arxiv.org/abs/1303.3997) and BWA (9), which are commonly used short read aligners for constructing a scaffolding graph (3). BOWTIE2 has two alignment modes, local and global, leading to the versions BOWTIE2 LOCAL and BOWTIE2 GLOBAL respectively. It is important to emphasise that FAST-SG will never be used to align reads to a finished reference genome. We performed this benchmark because it has been extensively used to demonstrate the performance of short read aligners and allows a more natural comparison at the read level between aligners and FAST-SG.

**Table 2.**
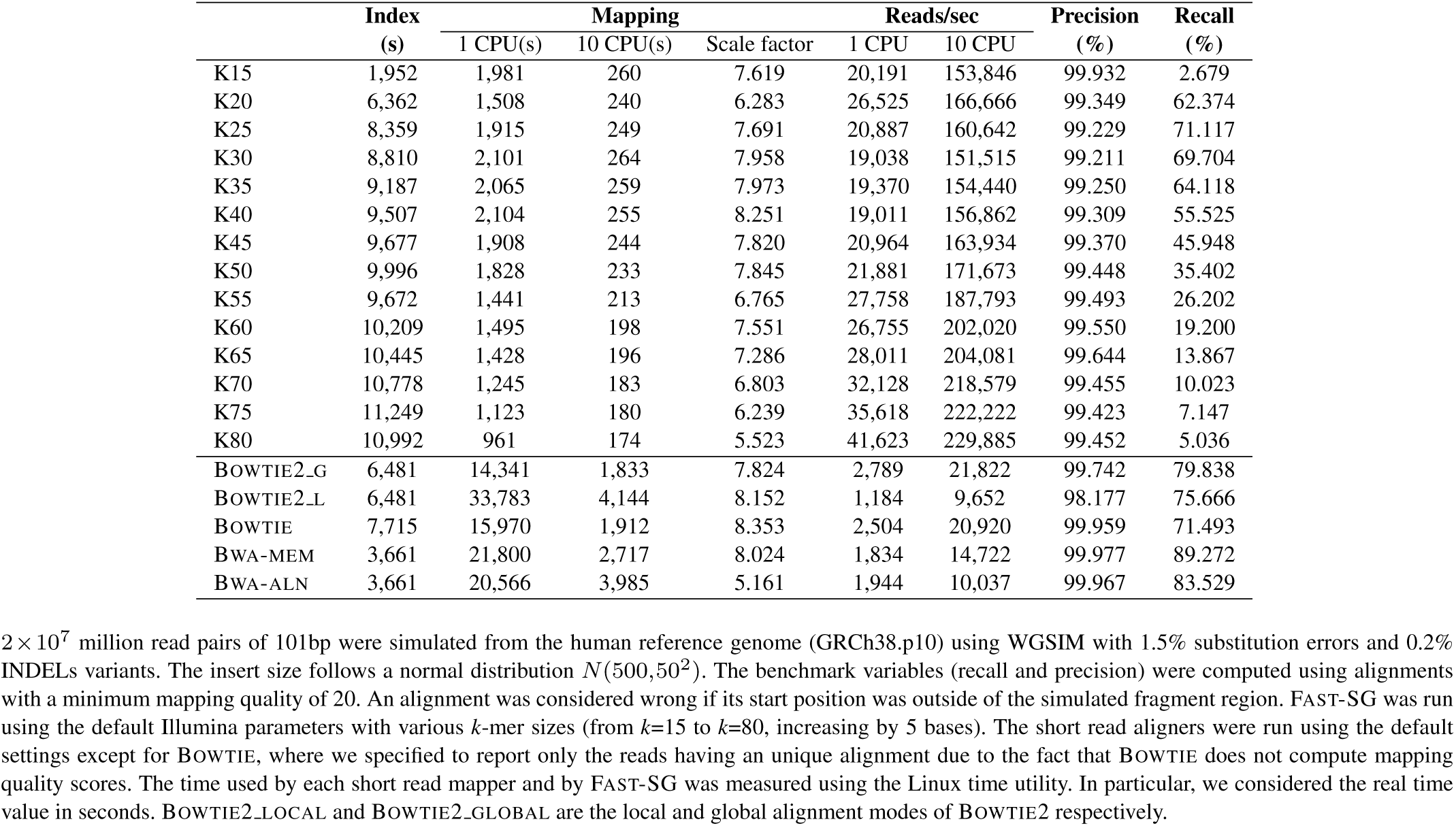
Illumina alignment benchmark.

Our results show that BWA-MEM is the leading tool in terms of Precision. FAST-SG is superior to BOWTIE2 LOCAL for any *k*-mer size and is comparable to BOWTIE2 GLOBAL. In terms of speed, FAST-SG performs the best. Indeed, it is between 7X to 14X times faster (depending on the *k*-mer size) than the next fastest program, which is BOWTIE2 GLOBAL. Moreover, the scale factor of FAST-SG when multiple CPUs are used is comparable to the one obtained by the short read aligners that were evaluated so that the ratio in terms of speed among the different aligners remains the same. Notice that FAST-SG can be the slowest tool when constructing the index depending on the *k*-mer size used. In the worst case, it is however only 3X times slower than BWA, which is the fastest tool. With respect to recall, BWA-MEM is the leading tool. The recall of FAST-SG depends on the *k*-mer size used (Table 2, Figure 1b). Larger *k*-mer values (k*>*50) decrease the recall of FAST-SG due to sequencing errors and read length. For *k* between 25 and 30, which corresponds to optimal *k*-mer sizes, the recall of FAST-SG is comparable to the one of BOWTIE (Table 2).

### Comparison of FAST-SG with the short read aligners commonly used for constructing a scaffolding graph

A more balanced way of evaluating FAST-SG is to compare its performance to the one of other commonly used short read alignment tools for scaffolding graph construction in terms of the scaffolding results. Indeed, Hunt *et al.* (3) demonstrated that the quality of the latter is highly dependent on the short read aligner used and that precision is more important than maximizing the number of reads aligned to the contigs.

Such comparison was done on four real test cases (Table 1, Supplementary Material 3) using two well established scaffolders, namely OPERA-LG (11) and BESST2 (12), and two more recently published ones, namely SCAFFMATCH (13) and BOSS (14). All scaffolders have different algorithms to select optimal paths from the scaffolding graph. They support the SAM/BAM format and were run with identical commands overall (Supplementary Material 3).

In relation to the number of paired reads mapped (Supplementary Figure S1), FAST-SG aligned on average more pairs than BOWTIE or BWA, and was comparable to BOWTIE2 GLOBAL. It however aligns less pairs than BOWTIE2 LOCAL or BWA-MEM. From the number of paired reads aligned across the four test cases, we notice that the behavior of FAST-SG depends on the *k*-mer size chosen. With larger sizes, FAST-SG resembles global methods and with shorter sizes, it is closer to local methods (Supplementary Figure S1).

The average contig read-coverage statistics which is used to tag the repeated contigs before scaffolding (2) was extracted from the results of OPERA-LG. This statistics was employed to compute a pairwise Pearson correlation to determine the linear relationship between the short read aligners and FAST-SG (Supplementary Figure S2). We observe that the average contig read-coverage computed from the FAST-SG alignments correlated more on average with BOWTIE (*x*=0.933), BWA (*x*=0.905) and BOWTIE2 GLOBAL (*x*=0.814) than with BWA-MEM (*x*=0.772) or BOWTIE2 LOCAL (*x*=0.725) on the datasets of *S. aureus*, *R. sphaeroides* and *P. falciparum* (Supplementary Figure S2).

To evaluate the scaffolding results (Methods section, Supplementary Material 3), the number of correct and erroneous joins were computed in each test case using the scripts provided in Hunt *et al.* (3). Moreover, the F-Score metric (Methods section) was employed to summarise in a single statistics the performance of each scaffolder when using FAST-SG or the short read aligners.

The results of the four test cases in terms of F-score and error rate are illustrated in Figure 2 and detailed in Supplementary Tables S1 to S4. For almost all the test cases and scaffolding tools, FAST-SG reached the largest F-score (Figure 2) for some *k*-mer value. Moreover, FAST-SG had a superior average performance in terms of F-score in relation to the four scaffolders tested in 2 out of the 5 datasets (Figure 2, vertical lines) and allowed the scaffolding tools to obtain more accurate scaffolding results in 4 out of the 5 datasets (Figure 2, vertical lines).

**Figure 2.**
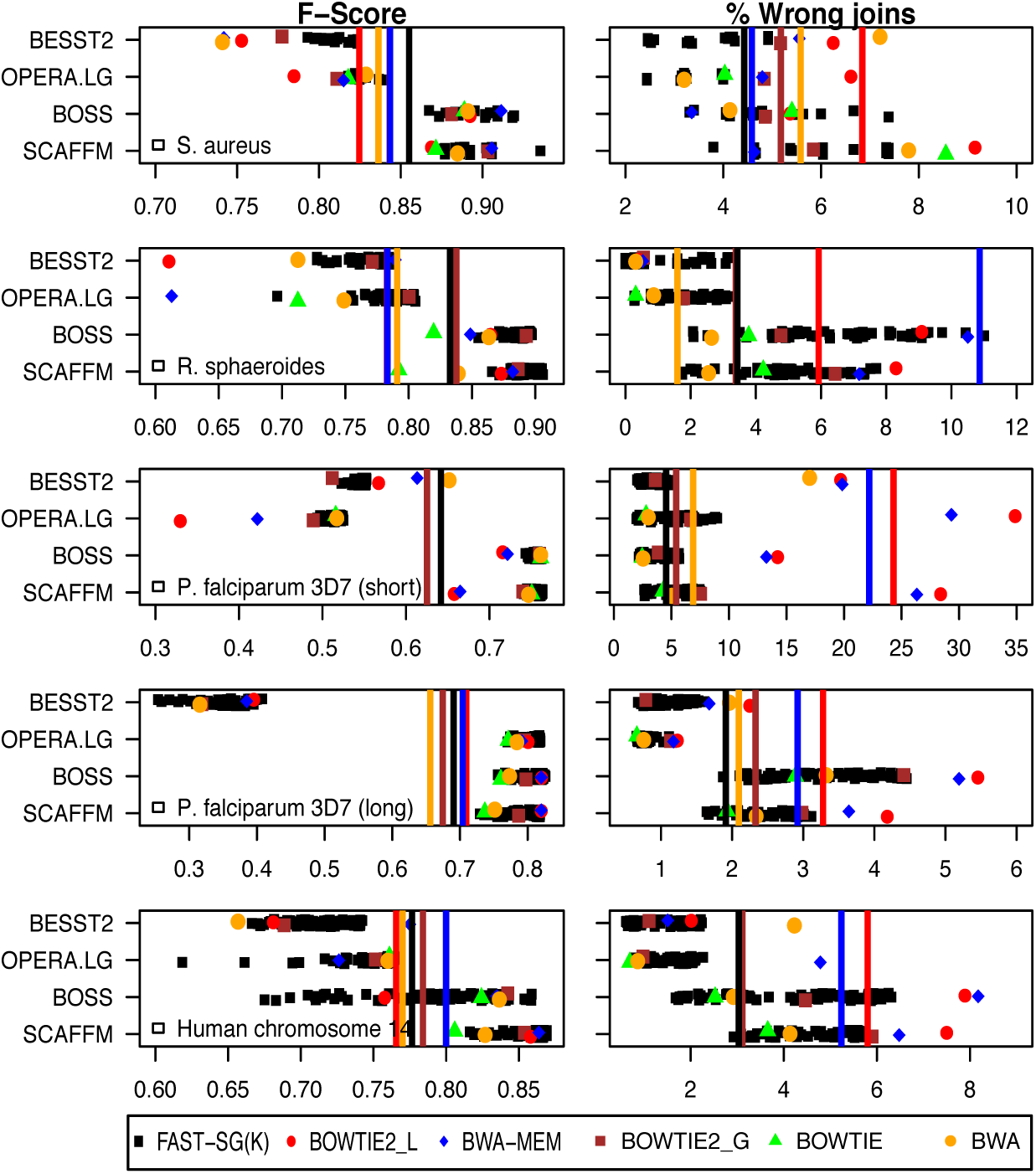
Illumina scaffolding benchmark. Four real datasets (Table 1), five Illumina libraries and four scaffolding tools were used to assess the performance of F_AST_-SG and the short read aligners for building the scaffolding graph by means of an F-score metric and percentage of wrong joins (Methods Section, and Supplementary Material 3). F_AST_-SG was run with various *k*-mer sizes in the range of *k*=12-28, *k*=12-70, *k*=15-66 and *k*=15-80 for *Staphylococcus aureus*, *Rhodobacter sphaeroides*, *Plasmodium falciparum* and the human chromosome 14, respectively. Short read aligners were run with the wrapper or instructions provided by the scaffolding tools when possible, or using the default parameters. Single data points provide the F-Score and error rate for each combination of scaffolding tool and aligner in each dataset. The vertical lines show for each dataset the average F-score or Error rate values obtained by each of the short read aligners or F_AST_-SG together with the four scaffolding tools. Vertical lines for B_OWTIE_ were not plotted since it cannot be used with B_ESST_2. For the *Plasmodium falciparum* (short) dataset, the average F-Score (vertical lines) were omitted for B_WA_, B_WA_-_MEM_ and B_OWTIE_2 L_OCAL_ due to a poor performance (High error rate). The commands used for the aligners and scaffolding tools are detailed in the Supplementary Material 3.

The low GC content genome of *Plasmodium falciparum* proved to be particularly challenging to the scaffolders using local alignment methods (namely BWA-MEM or BOWTIE2 LOCAL). These indeed tended to produce several wrong joins (Figure 2), indicating that the local alignment methods are not an appropriate choice for scaffolding this genome. A possible explanation for the poor performance observed in this particular case is that the local alignment methods mapped 10% more reads than the global ones and than FAST-SG (Supplementary Figure S1), but there is a low correlation in the average contig read-coverage between the local alignment methods and FAST-SG (Supplementary Figure S2), suggesting many wrong mappings in the extra 10% aligned reads.

Concerning the scaffolding tools, BOSS and SCAFFMATCH reached higher F-score values than OPERA-LG and BESST2 (Figure 2). However, they tended to produce more scaffolding errors (Figure 2). Additionally, it is important to observe that fragmented Illumina assemblies (Table 1) were used to maximize the number of gaps to spam and better assess the influence of the short read aligners in the scaffolding process.

In conclusion, over the four test cases and four scaffolders benchmarked, FAST-SG consistently reached better scaffolding results than the short read aligners evaluated and may be considered as a reliable tool for constructing a scaffolding graph from short reads.

### Building synthetic read pairs libraries from long reads

Despite the high per-base error rate of the long reads technologies, the long-range information encoded in a long read has proven to be highly accurate. On the other hand, current experimental protocols to produce long-range mate-pair libraries using short read technologies are time-consuming and expensive (22, 23). Moreover, library contamination occurs when the circularization step fails during construction, resulting in mate-pairs with short insert size and in the wrong orientation (12). Extracting synthetic mate-pair libraries directly from long reads could improve the performance of the current short read scaffolders and replace the need for sequencing multiple mate-pair libraries for scaffolding.

To demonstrate the utility of FAST-SG to create synthetic mate-pair libraries from long reads, we collected the latest chemistry data sequenced with the Oxford Nanopore (1D reads sequenced on R9.2 flow cells) and Pacific Biosciences (Sequel System) technologies (respectively denoted by ONT and PacBio from now on) for the genome of *Escherichia coli* K12 (Table 1, Supplementary Table S3). The long reads were error-corrected using Illumina reads (Table 1, Supplementary Material 4) with LORDEC (30), a hybrid error-correction method.

FAST-SG was used to generate synthetic mate-pair libraries in the range of 0.5-8kb from the corrected and uncorrected long reads using a *k*-mer size of 15, at which 98% of the *k*-mers are unique in the reference *E. coli* K12 genome. Synthetic mate-pair reads were aligned to an Illumina assembly of *E. coli K12* (Table 1). Near perfect synthetic mate-pair libraries were obtained with a low percentage of outliers (<9.85%) for all insert sizes (Figure 3). Moreover, the hybrid error-correction reduced the standard deviation and allowed the average insert size to get close to the specified size of each synthetic library. However, the hybrid error-correction increased the number of outliers in both technologies (Figure 3). The observed average insert size (Figure 3) in the synthetic libraries from ONT are slightly higher than the observed ones in PacBio, thus reflecting the nature of the error of each long read technology, which are deletions for ONT (4) and substitutions for PacBio (4).

**Figure 3.**
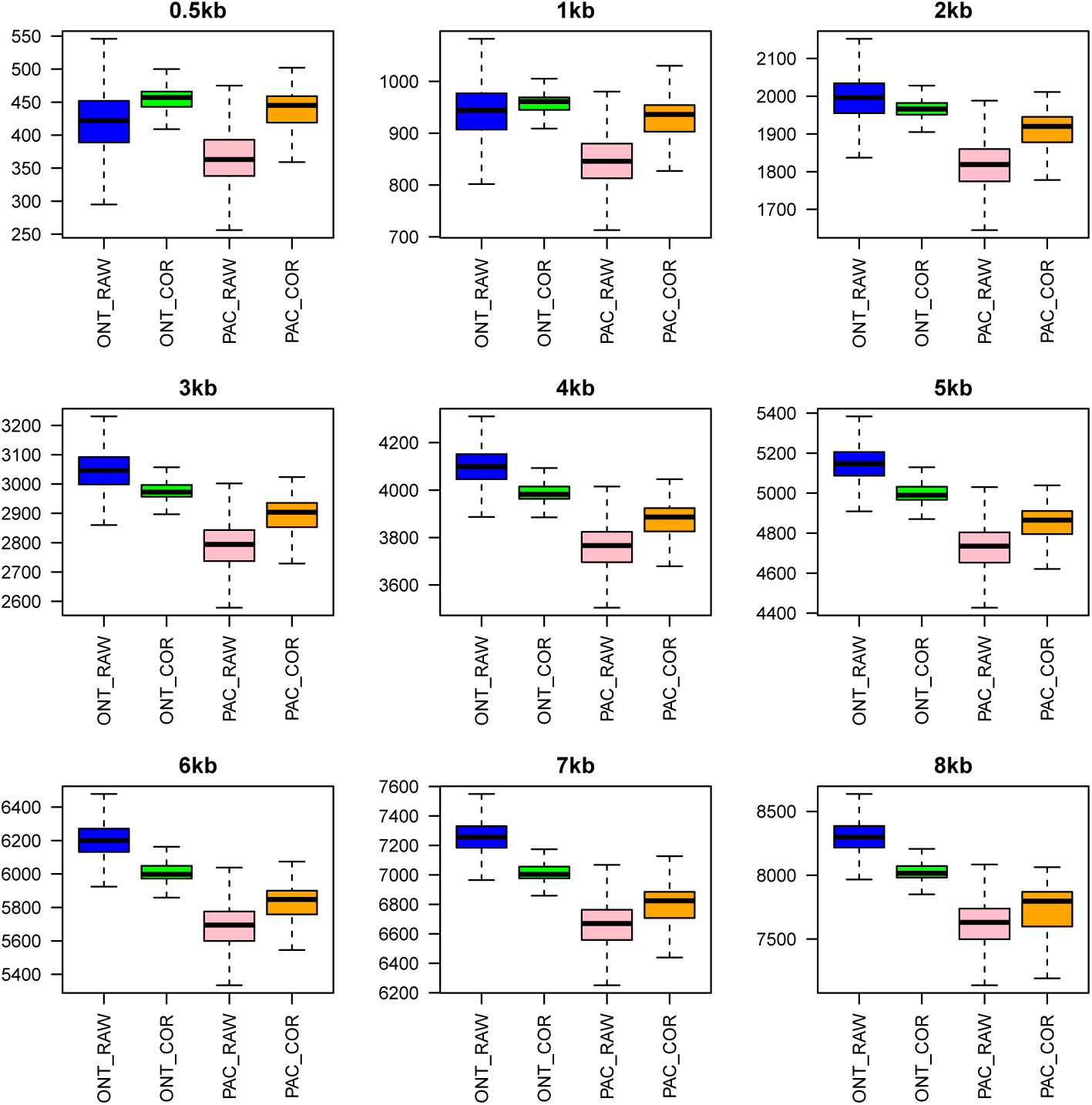
Boxplots of the insert size distribution observed for each synthetic library in the genome of *Escherichia coli* K12. The boxplots were drawn extracting from the F_AST_-SG alignments a minimum of 5,000 insert sizes from the mate-pair reads mapped within contigs for each combination of synthetic library and long read technology. The percentage of outliers detected in the raw ONT reads ranged from a minimum of 0.37% (0.5kb) to a maximum of 4.24% (8kb), while for raw PacBio it ranged from a minimum of 0.25% (0.5kb) to a maximum of 9.85% (8kb). The number of outliers increased with the error-correction for both long read technologies, reaching an average of 9.32% (std 1.73%) and 8.32% (std 3.74%) for the ONT and PacBio reads, respectively. The boxplots were drawn excluding outliers.

On this dataset, we computed the recall achieved by FAST-SG at the levels of the *k*-mers and of the synthetic mate-pair reads (the length of the forward and reverse reads equals 200 base pairs) for each long read technology, from either raw or corrected reads (Supplementary Table S8). At the *k*-mer level, FAST-SG has a recall of 8.3% and 5.05% for the uncorrected reads of ONT and PacBio, respectively. The hybrid error-correction increased the *k*-mer recall by 10% for both long read technologies. At the synthetic mate-pair read level, we observed a recall of 49.42% and 31.65% for the raw ONT and raw PacBio reads, respectively. The hybrid error-correction increases the synthetic mate-pair read recall for ONT to 75.12% and for PacBio to 65.02%. We observed that FAST-SG is more effective aligning synthetic mate-pair reads from raw ONT than from raw PacBio reads. We expect that this is due to the nature of the ONT errors (major deletions) as FAST-SG is designed to deal with short indels. Despite the low *k*-mer recall, FAST-SG achieved a decent synthetic mate-pair read recall on this dataset from both long read technologies, and extracted near perfect synthetic mate-pair libraries. The synthetic mate-pair libraries can be used as input to a short read scaffolder to generate scaffold sequences through a combination of short and long read technologies.

### Comparison of FAST-SG coupled with short read scaffolders against LINKS

We compared the results obtained by FAST-SG coupled with the short read scaffolders against LINKS (35), which is a scaffolder specifically designed to extract paired *k*-mers from long reads and employ them to join contigs.

FAST-SG and LINKS were applied with default parameters (*k*-mer of size 15) to create the synthetic mate-pair libraries in the range of 0.5kb to 8kb using as input the uncorrected long reads and Illumina assemblies available for both species (Table 1, Supplementary Material 4). Since LINKS performs better with high long read coverage (35), we subsampled 50X and 30X of coverage from *Escherichia coli* K12 and *Saccharomyces cerevisiae* W303, respectively.

FAST-SG is two times faster than LINKS and requires two orders of magnitude less memory to extract linking information from the long reads (Supplementary Table S9). The percentages of linked pairs extracted by both methods is comparable (with FAST-SG being slightly superior) and as expected, the percentage of linked pairs increases as a function of the insert size length for both long read technologies (Supplementary Table S10).

A more informative comparison involved assessing the quality of the scaffolds (3) produced by LINKS on one hand, and on the other, by the short read scaffolders coupled with FAST-SG. Based on the F-Score values (Methods section), the short read scaffolders using FAST-SG reached better or comparable results than LINKS (Figure 4). Moreover, LINKS produced more scaffolding errors in two out of the three datasets tested (Supplementary Table S11). With respect to the *E. coli* dataset, the scaffolding errors (Methods section) made by the short read scaffolders using FAST-SG (Figure 4) were related to the gap size estimation (type error 4), orientation (type errors 1 and 5), and relocation (type errors 8 and 12). The major source of errors in the scaffolds produced by LINKS was of type 5. This measures the correct orientation and distance between pairs of contigs (Pie chart in Figure 4). On the *S. cerevisiae* W303 dataset, the major source of scaffolding errors was translocation (type error 2) for both methods. However, LINKS has almost double the number of scaffolding errors compared to FAST-SG coupled with OPERA-LG or BOSS on this dataset (Supplementary Table S11).

**Figure 4.**
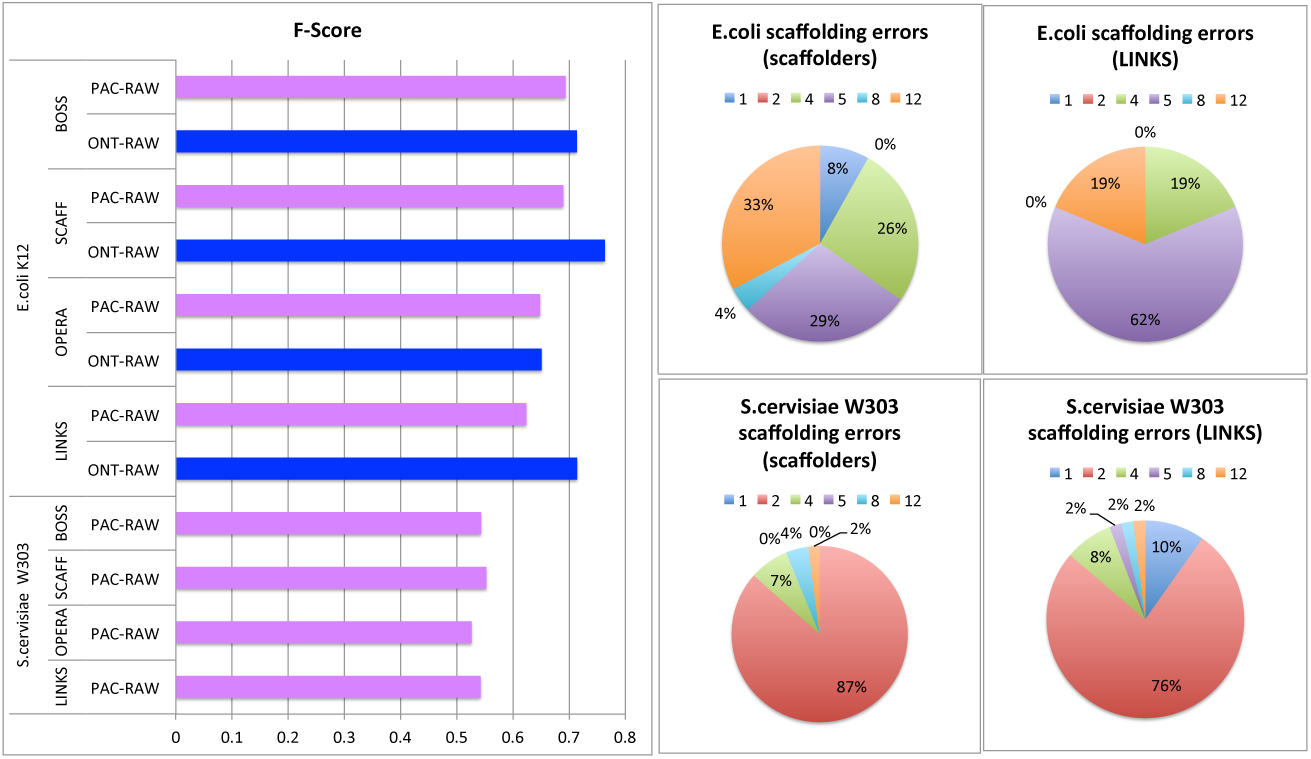
Synthetic libraries scaffolding benchmark. The F-Score (Methods Section) was computed with the scripts provided by Hunt *et al*. (3) on the scaffold sequences produced by each scaffolding tool. The pie charts display the scaffolding error details for L_INKS_ and for the short read scaffolders fed with the F_AST_- SG alignments for both *E. coli* K12 and *S. cerevisiae* W303. The definition of the scaffolding errors (numbers in the pie charts) are provided in the Methods section.

Overall, the performance of the short read scaffolders coupled with FAST-SG was superior or comparable to LINKS, a scaffolder specifically designed for long reads. FAST-SG thus allows the conversion of tools designed for short read scaffolding into a long read scaffolder in a fast and modular way. It is important to notice also that the scaffolding errors observed here can be further reduced because fragmented Illumina assemblies (Table 1) were used in order to maximize the possibility of the scaffolders to make joins.

### Using FAST-SG to perform the hybrid assembly of *Arabidopsis thaliana* (Ler-0)

In this experiment, we demonstrate the utility of FAST-SG as part of a hybrid assembly pipeline. Additionally, we explore the long read coverage required to obtain long range scaffold sequences comparable to CANU (4), a state-of-the-art *de novo* long read assembly pipeline.

Briefly, the hybrid assembly using FAST-SG proceeded as follows. In a first step, a single Illumina library (Table 1) covering 100X the *Arabidopsis thaliana* (Ler-0) genome was assembled using DISCOVAR_*denovo*_ (33), which is one of the best tools for assembling a single Illumina fragment (pair-end) library. The resulting assembly contained a total of 2,384 scaffolds with a N50 of 320kb and a total size of 119Mb (Table 3). The DISCOVAR_*denovo*_ assembly took 6.6 hours on 20 CPUs. In a second step, a total of 50X of PacBio reads (P5-C3) were error-corrected (Table 1), with the same Illumina reads used for the *de novo* assembly, using LORDEC. LORDEC took 14.2 hours on 20 CPUs. In a third step, the error-corrected long reads were randomly subsampled with a coverage between 5X to 50X, and FAST-SG (using 21-mers) was used to create 12 synthetic mate-pair libraries in the range of 1kb to 20kb for each subsample. The total number of mate-pair reads aligned at each coverage value ranged from 11.85 to 104.99 million for 5X to 50X, respectively (Supplementary Table 12). On average, 7.2% of the synthetic mate-pair reads aligned by FAST-SG were linking (i.e. connecting two different contigs) in each subsample. Moreover, a near perfect insert size distribution and a low percentage of outliers were observed for each synthetic library (Supplementary Figure S3). FAST-SG took 2.15 hours on 20 CPUs to process the whole dataset. Finally, OPERA-LG, BOSS and SCAFFMATCH were fed with the FAST-SG alignments to produce the scaffold sequences (Table 3). All short read scaffolders generated scaffold sequences in at most half an hour (OPERA-LG 22min, BOSS 24m and SCAFFMATCH 30min) using a single CPU.

**Table 3.** Hybrid and *de novo* assemblies of *Arabidopsis thaliana* (Ler-0)

| Number<br>Scaffolds | Max | N50 | Size<br>(Mb) | Fold | LRC | Scaffolder /<br>Assembler | BreakPoints |  |  | 1-to-1<br>identity | % Ref<br>covered |
| --- | --- | --- | --- | --- | --- | --- | --- | --- | --- | --- | --- |
|  |  |  |  |  |  |  | Number | Bases (Mb) | % Error |  |  |
| 2,384 | 1,551,485 | 320,571 | 119.45 | 1.00 | - | DISCOVAR | 91 | 0.48 | 0.49 | 99.07 | 82.044 |
| 1,577 | 5,305,497 | 1,076,408 | 120.05 | 3.36 | 5X | OPERA-LG | 174 | 0.978 | 1.00 | 99.07 | 82.054 |
| 1,368 | 9,953,317 | 2,475,756 | 120.26 | 7.72 | 10X | OPERA-LG | 202 | 1.197 | 1.22 | 99.07 | 82.047 |
| 1,249 | 16,906,870 | 4,165,132 | 120.32 | 12.99 | 15X | OPERA-LG | 206 | 1.237 | 1.26 | 99.07 | 82.052 |
| 1,179 | 18,032,662 | 4,941,257 | 120.41 | 15.41 | 20X | OPERA-LG | 218 | 1.588 | 1.62 | 99.07 | 82.060 |
| 1,103 | 14,710,653 | 4,756,724 | 120.43 | 14.84 | 30X | OPERA-LG | 227 | 1.728 | 1.76 | 99.07 | 82.055 |
| 1,049 | 10,003,725 | 4,667,601 | 120.41 | 14.56 | 50X | OPERA-LG | 230 | 1.732 | 1.76 | 99.07 | 82.060 |
| 1,345 | 8,867,374 | 1,632,787 | 120.40 | 5.09 | 5X | SCAFFM | 195 | 1.620 | 1.65 | 99.07 | 82.058 |
| 1,143 | 8,867,059 | 5,142,417 | 120.65 | 16.04 | 10X | SCAFFM | 203 | 1.319 | 1.34 | 99.07 | 82.045 |
| 1,072 | 11,814,750 | 6,165,459 | 120.73 | 19.23 | 15X | SCAFFM | 205 | 1.330 | 1.36 | 99.07 | 82.045 |
| 1,020 | 11,873,221 | 6,221,109 | 120.80 | 19.41 | 20X | SCAFFM | 207 | 1.477 | 1.50 | 99.07 | 82.039 |
| 958 | 13,946,812 | 7,073,179 | 120.90 | 22.06 | 30X | SCAFFM | 209 | 1.651 | 1.68 | 99.07 | 82.042 |
| 923 | 13,957,620 | 6,292,557 | 120.85 | 19.63 | 50X | SCAFFM | 210 | 1.712 | 1.74 | 99.07 | 82.041 |
| 1,593 | 5,296,335 | 1,037,785 | 119.96 | 3.24 | 5X | BOSS | 179 | 1.171 | 1.19 | 99.07 | 82.061 |
| 1,371 | 13,608,688 | 2,554,739 | 120.17 | 7.97 | 10X | BOSS | 200 | 1.335 | 1.36 | 99.07 | 82.054 |
| 1,239 | 13,643,115 | 2,829,628 | 120.22 | 8.83 | 15X | BOSS | 207 | 1.189 | 1.21 | 99.07 | 82.061 |
| 1,173 | 7,977,908 | 3,005,451 | 120.23 | 9.38 | 20X | BOSS | 212 | 1.564 | 1.59 | 99.07 | 82.060 |
| 1,093 | 9,004,636 | 2,974,378 | 120.28 | 9.28 | 30X | BOSS | 219 | 1.575 | 1.60 | 99.07 | 82.057 |
| 1,031 | 11,011,921 | 3,179,270 | 120.29 | 9.92 | 50X | BOSS | 229 | 2.162 | 2.20 | 99.07 | 82.050 |
| 1,439 | 447,211 | 80,063 | 89.84 | - | 10X | CANU | 107 | 0.675 | 1.10 | 98.19 | 51.188 |
| 259 | 4,542,617 | 1,170,676 | 118.25 | - | 20X | CANU-P | 201 | 0.969 | 0.99 | 99.06 | 81.907 |
| 258 | 4,543,625 | 1,170,942 | 118.31 | - | 20X | CANU-Q | 183 | 0.831 | 0.85 | 99.02 | 81.808 |
| 259 | 4,535,400 | 1,168,180 | 118.05 | - | 20X | CANU | 185 | 1.030 | 1.09 | 98.82 | 78.874 |
| 119 | 15,152,700 | 6,219,401 | 120.67 | - | 50X | CANU | 219 | 1.766 | 1.79 | 99.02 | 82.565 |
| 88 | 15,945,651 | 8,307,845 | 121.45 | - | 150X | CANU | 215 | 1.935 | 1.95 | 99.06 | 82.938 |

The hybrid and the CANU assemblies available were structurally validated by a whole genome alignment against the reference

*Arabidopsis thaliana* TAIR10 genome (Table 3, Methods section, Supplementary Material 4).

As can be seen from Table 3, all hybrid assembly pipelines were able to produce long-range scaffolds (N50 *>* 1Mb) with a high coverage of the reference genome, low number of errors (<2.2%), low amount of sequence gaps (1.45Mb as maximum), and with an identity higher than any CANU assembly. All hybrid assemblies at 5X of coverage reached a N50 scaffold size comparable to the contig N50 obtained by a polished CANU assembly requiring 20X of coverage and 100X of Illumina reads (Table 1). Additionally, all hybrid assembly pipelines seemed to plateau after 30X of long read coverage as was previously observed on this dataset (4). However, SCAFFMATCH, the most aggressive scaffolder tested, at 10X-30X of coverage produced accurate scaffolds having an N50 comparable to the CANU assemblies requiring 50X or 150X of coverage (Table 3).

All assemblies of *Arabidopsis thaliana* (Ler-0) were comparable in terms of the number and amount of sequences involved in structural errors (Table 3). Moreover, the major source of structural errors observed in both assembly strategies were mainly relocations, which explain more than 50% of the amount of sequences involved in miss-assemblies (Supplementary Figure S5).

Overall, we demonstrated that hybrid assemblies at low long read coverage were comparable in terms of continuity, completeness and accuracy to the assemblies obtained by CANU, which is considered a state-of-the-art *de novo* long read assembly pipeline. Indeed, hybrid assembly allowed cheaper (low long read coverage and a single Illumina library) and faster reconstruction of the *Arabidopsis thaliana* (Ler-0) genome.

### Using FAST-SG to perform the hybrid assembly of a diploid human genome (NA12878)

An ultimate benchmark for any assembly method or sequencing technology is to assemble a complete human genome (4, 19, 21, 31, 36). We performed a hybrid assembly of the Utah/Ceph NA12878 human diploid genome using a low coverage (5X) of ultra-long Nanopore reads (Table 1, BioRxiv: https://doi.org/10.1101/128835), a DISCOVAR_*denovo*_ assembly built from 50X of 250bp Illumina reads (Table 1, (33)), FAST-SG and SCAFFMATCH (13).

FAST-SG (using 22-mers) was run to create 20 synthetic mate-pair libraries in the range of 2kb-180kb using as input a total of 1.4 million uncorrected Nanopore reads (N50 64.75Kb, Table 1, Supplementary Table S6), which have a total size of 23.11Gb and cover about 7X of the human genome. A total of 455.9 million synthetic mate-pair reads (11.15% linking contigs, Supplementary Table 13) were aligned to the DISCOVAR_*denovo*_ assembly, with a near perfect distribution of insert sizes and a low percentage of outliers observed (Supplementary Figure S4). FAST-SG required 8 hours using 20 CPUs to complete the task and used a maximum of 25Gb of memory. SCAFFMATCH was then fed with the alignments of FAST-SG and took 5.18 hours using a single CPU with a peak memory of 30.87Gb to generate the scaffold sequences. The resulting hybrid assembly is referred here to as the DFS (DISCOVAR_*denovo*_ +FAST-SG+SCAFFMATCH) assembly.

We evaluated the accuracy of the DFS assembly together with the public assemblies of NA12878 that were built using CANU (BioRxiv: https://doi.org/10.1101/128835), MASURCA (31), 10X genomics (21) and DOVETAIL genomics (19) by means of whole genome alignments against the complete human reference genome (Table 4, Methods section).

**Table 4.**
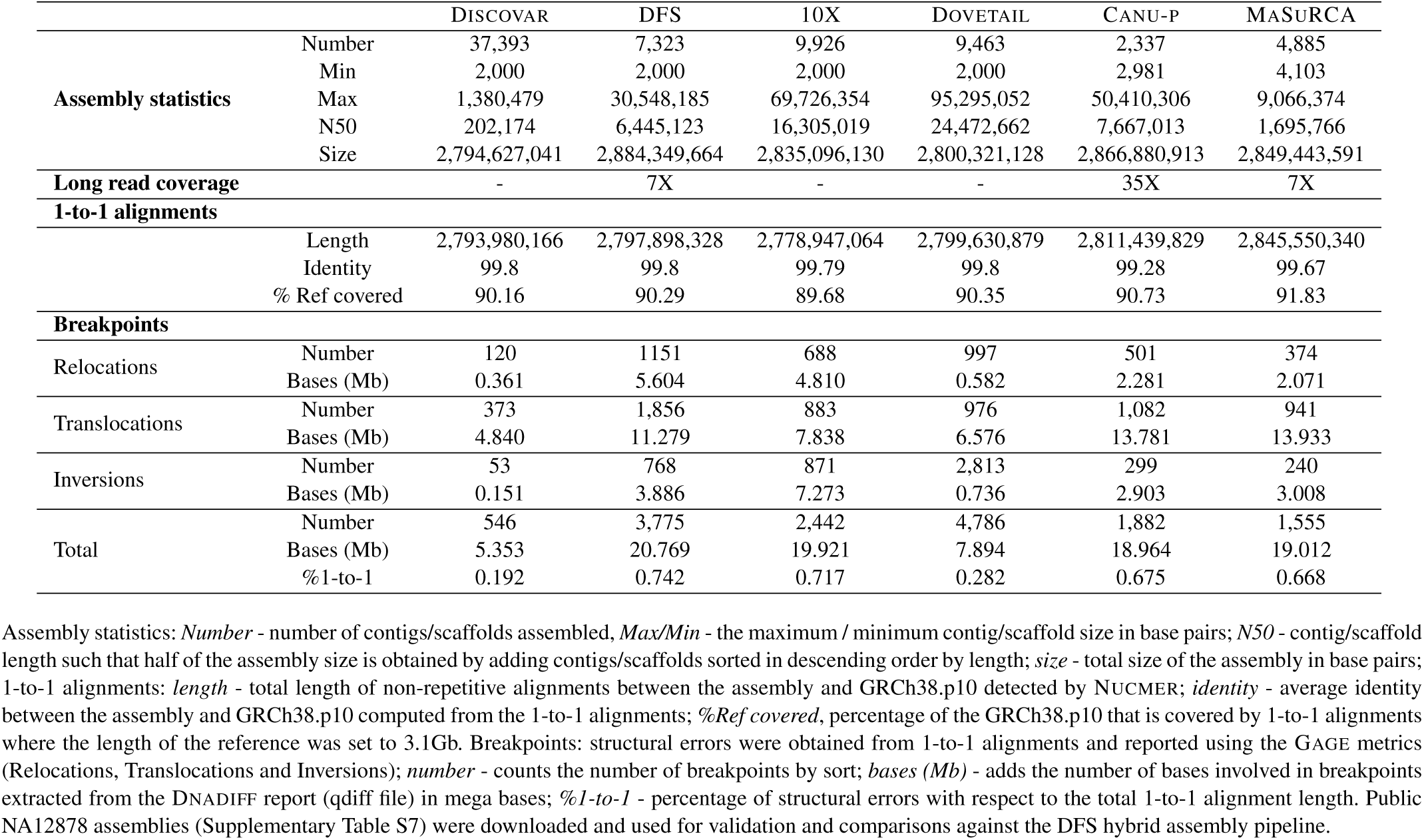
Hybrid and *de novo* assemblies of NA12878

|  |  | DISCOVAR | DFS | 10X | DOVETAIL | CANU-P | MASURCA |
| --- | --- | --- | --- | --- | --- | --- | --- |
| Assembly statistics | Number | 37,393 | 7,323 | 9,926 | 9,463 | 2,337 | 4,885 |
|  | Min | 2,000 | 2,000 | 2,000 | 2,000 | 2,981 | 4,103 |
|  | Max | 1,380,479 | 30,548,185 | 69,726,354 | 95,295,052 | 50,410,306 | 9,066,374 |
|  | N50 | 202,174 | 6,445,123 | 16,305,019 | 24,472,662 | 7,667,013 | 1,695,766 |
|  | Size | 2,794,627,041 | 2,884,349,664 | 2,835,096,130 | 2,800,321,128 | 2,866,880,913 | 2,849,443,591 |
| Long read coverage |  | - | 7X | - | - | 35X | 7X |
| 1-to-1 alignments |  |  |  |  |  |  |  |
|  | Length | 2,793,980,166 | 2,797,898,328 | 2,778,947,064 | 2,799,630,879 | 2,811,439,829 | 2,845,550,340 |
|  | Identity | 99.8 | 99.8 | 99.79 | 99.8 | 99.28 | 99.67 |
|  | % Ref covered | 90.16 | 90.29 | 89.68 | 90.35 | 90.73 | 91.83 |
| Breakpoints |  |  |  |  |  |  |  |
| Relocations | Number | 120 | 1151 | 688 | 997 | 501 | 374 |
|  | Bases (Mb) | 0.361 | 5.604 | 4.810 | 0.582 | 2.281 | 2.071 |
| Translocations | Number | 373 | 1,856 | 883 | 976 | 1,082 | 941 |
|  | Bases (Mb) | 4.840 | 11.279 | 7.838 | 6.576 | 13.781 | 13.933 |
| Inversions | Number | 53 | 768 | 871 | 2,813 | 299 | 240 |
|  | Bases (Mb) | 0.151 | 3.886 | 7.273 | 0.736 | 2.903 | 3.008 |
| Total | Number | 546 | 3,775 | 2,442 | 4,786 | 1,882 | 1,555 |
|  | Bases (Mb) | 5.353 | 20.769 | 19.921 | 7.894 | 18.964 | 19.012 |
|  | %1-to-1 | 0.192 | 0.742 | 0.717 | 0.282 | 0.675 | 0.668 |

In terms of continuity (N50, Table 4), the DFS assembly is more than 4X larger than a MASURCA hybrid assembly built with the same long read dataset and 100X of Illumina reads (http://masurca.blogspot.cl/2017/06/masurca-assembly-of-na12878-low.html). Moreover, it is comparable to a polished CANU assembly built with 35X of long read coverage (BioRxiv: https://doi.org/10.1101/128835). DOVETAIL genomics and 10X genomics reached larger N50 scaffolds (Table 4), which are 2.5X and 3.7X larger than the DFS assembly, respectively. All assemblies are comparable in terms of size, 1-to-1 alignment length and coverage of the reference genome (Table 4).

In terms of identity (Table 4), DOVETAIL genomics and DFS are the leading pipelines. DOVETAIL genomics and DFS both use the DISCOVAR_*denovo*_ assembly as input for scaffolding. Both tools maintain the high identity of the DISCOVAR_*denovo*_ assembly because contig bases are not changed in the scaffolding process.

As concerns the structural errors, all assembly pipelines are highly accurate with less than 1% of the total 1-to-1 alignment length involved in such errors (Table 4, Supplementary Figure S6). Moreover, translocation is the structural error that accumulates the greatest amount of miss-assembled bases on all assembly pipelines (Table 4). A more detailed inspection of the 1-to-1 alignments revealed that DFS, 10X genomics and DOVETAIL genomics tend to skip the short contigs (Supplementary Table S14), which is a known problem of scaffolding tools (3). However, more complex miss-assemblies involving several structural errors were observed in the chimeric contigs assembled by CANU and MASURCA (Supplementary Table S15).

In terms of speed, the whole DFS pipeline (933 CPU hours) was 22X times faster than MASURCA (21.000 CPU hours; personal communication), 162X times faster than CANU (151.000 CPU hours BioRxiv: https://doi.org/10.1101/128835), and comparable to 10X genomics and DOVETAIL genomics.

Finally, we call attention to the fact that the hybrid assembly solution which we propose (using 14 flow cells and 50X of 250bp PE reads sequenced on Hiseq2500) is approximately 3 times cheaper than the CANU solution (using 53 flow cells and 50X of Illumina).

In summary, we demonstrated in this experiment that the DFS hybrid assembly pipeline produced an accurate and long-range reconstruction of a diploid human genome that was faster and cheaper than the current state-of-the-art long read assembly pipelines.

## Discussion

We introduced in this paper a new method, FAST-SG, that enables to construct a scaffolding graph from either short or long reads, allowing for an accurate construction of the scaffold sequences as well as for software reuse.

We started by showing that on standard Illumina benchmarks, FAST-SG is faster than the current state-of-the-art short read aligners and that better results are achieved by the scaffolding tools when they are coupled with FAST-SG.

We then showed that near perfect synthetic libraries are obtained with FAST-SG from either corrected or uncorrected PacBio and Nanopore long reads. The insert size is restricted to the actual long read size, but FAST-SG is able, using ultra-long Nanopore reads, to extract synthetic libraries of even Bacterial Artificial Chromosome clone sizes having insert size of 150kb-180kb. Those kinds of libraries were crucial to reach the high continuity of the current human reference genome (36). An estimation of the gap size with the existing long-range mate-pair technologies (10X genomics and DOVETAIL genomics) is more challenging than with the synthetic libraries due to the fact that in such technologies, the linking information comes from a range of insert-sizes and the relative orientation of the read pairs may not be known (DOVETAIL genomics).

Clearly, the synthetic libraries eliminate the bottleneck of sequencing a combination of mate-pair libraries, which were typically required to obtain long-range assemblies (2, 22, 23). We further showed that short read scaffolders are able to produce accurate scaffolds when they are fed with the synthetic libraries extracted by FAST-SG, thus leading to results that are superior to or match those obtained by LINKS, a scaffolder specifically designed for hybrid long read scaffolding.

Finally, we demonstrated that FAST-SG in conjunction with efficient algorithms designed for Illumina data can be used to perform a full hybrid assembly of large genomes. The resulting assemblies are superior or comparable to the current state-of-the-art long read assembly pipelines. Furthermore, the modular hybrid pipelines are faster and cheaper in terms of computational resources and amount of long read coverage needed.

For further improvements of such hybrid assemblies, it would be possible to use the sequence between the synthetic pairs (“fake” gaps), either for assigning a new weight to the edges before scaffolding, or for placing the skipped contigs after scaffolding. An edge of the scaffolding graph can be re-weighted by computing the edit distance among the gap sequences and then eliminating the pairs having a large edit distance. EDLIB (37) is an efficient library that could be used to perform this task. The skipped contigs can be unambiguously placed by computing a consensus sequence of the scaffolding gaps, and then aligning the skipped contigs to the consensus gap sequence taking into account the lengths of the gap and of the skipped contig. The consensus of the sequence gaps can be computed in a faster way using the SPOA library, which implements a partial order alignment algorithm (38). These two improvements coupled with an appropriate ultra-long Nanopore read coverage (10X) could lead to an hybrid assembly pipeline that is superior to the current long-range mate-pair technologies where these improvements are not possible due to the fact that, in both technologies, the gap sequence between pairs is unknown. Clearly, improvement in the base accuracy of long reads will increase the recall of FAST-SG and thus impact positively on the hybrid assembly process. Notice however that read recall is less important because not all the sequenced reads are useful for scaffolding, and indeed we showed with the Illumina scaffolding benchmarks that the short read aligners with higher read recall produced the worst results. Additionally, FAST-SG was designed to enable constructing the scaffolding graph from uniquely mapped read pairs (FAST-SG index). It thus discards any repetitive sequence as they are not useful to build the scaffolding graph. Oxford Nanopore is a fast evolving technology and currently utilization of the new 1*D*^2^ chemistry or improvement in the base callers are two alternatives that could lead to an increased base accuracy of the ONT reads.

Overall, we believe that FAST-SG opens a door to achieve accurate hybrid long-range reconstructions of large genomes with low effort, high portability and low cost.

## Availability

FAST-SG is open source software under the MIT license available at https://github.com/adigenova/fast-sg.git.

## Funding

This work was supported by Basal program PFB03 and CONICYT PFCHA/BECA DOCTORADO NACIONAL 2014/FOLIO 21140124 granted to ADG. This research was partially supported by the supercomputing infrastructure of the NLHPC (ECM-02).

## Conflict of interest statement

None declared.

